# The assessment of single nucleotide polymorphisms in the ß-tubulin genes in human soil-transmitted helminths exposed to different pressure with benzimidazole drugs

**DOI:** 10.1101/2024.06.04.597280

**Authors:** Bruno Levecke, Nour Rashwan, Piet Cools, Marco Albonico, Shaali M Ame, Mio Ayana, Daniel Dana, Jennifer Keiser, Antonio Montresor, Zeleke Mekonnen, Sara Roose, Somphou Sayasone, Jozef Vercruysse, Jaco J Verweij, Johnny Vlaminck, Roger Prichard

## Abstract

**Background:** We aimed to gain insights into the role of known single nucleotide polymorphisms (SNPs) in codons 167, 198 and 200 of the ß-tubulin gene as markers for possible benzimidazole resistance in human soil-transmitted helminths (STHs; *Ascaris lumbricoides, Trichuris trichiura*, *Necator americanus* and *Ancylostsoma duodenale*).

**Methods:** Firstly, we determined the analytical performance of our PCR/pyrosequencing assays. Secondly, we applied them on stool samples collected during clinical trials in Ethiopia, Lao PDR, and Pemba Island (Tanzania) to assess any associations between the presence/ratio of mutant (MT): wild type (WT) SNPs and drug pressure history, individual drug response and time of sampling (baseline *vs.* follow-up sample).

**Principal findings:** Overall, the limit of blank of our in-house PCR/pyrosequencing assays to detect MT SNPs was non-zero (∼3.5%), and hence the limit of detection for MT SNPs was relatively high (2% – 7%). The assays systematically overestimated the true underlying ratio of MT:WT SNPS within sample, but we derived functions for more accurate estimates. The assays were more precise when the ratio MT:WT SNPs was high (>5%). No PCR amplicon was observed in 25% of the samples subjected to PCR. In the remaining samples, the presence of MT SNPs in codon 200 was detected in half of the analysed *Trichuris* samples, the proportion of the analysed samples containing MT SNPs did not exceed 14% for all other codons and STH species. Associations between drug pressure history, individual drug response and time of sampling, were not consistent across all codons and STHs.

**Conclusion:** We could not provide compelling evidence for the role of the known SNPs in the ß-tubulin gene as markers for benzimidazole resistance. Our study also highlights that there is a need to assess the diagnostic performance of any assays in order to readily interpret and compare results. Further research should therefore also focus on genes other than the ß-tubulin genes.

**Author summary:** Although large-sale deworming programs are reducing the morbidity caused by intestinal worms, widespread treatment of large populations for a long period of time may trigger drug resistance. An early detection of DNA mutations that may give rise to resistant worm population is therefore important. We evaluated the analytical performance of in-house assays to detect DNA mutations that are known to cause resistant intestinal worms of animals. Subsequently, we applied these assays on stool samples to verify (i) whether the mutations are more prevalent in areas were large proportions of children have been dewormed for a longer period, (ii) a poor individual drug response can be explained by higher frequency of the mutations. Our results indicate that comprehensive evaluation of the analytical performance of the genotyping tests was required to readily interpret the results. We did not find any compelling evidence that the presence of mutations was associated with either drug pressure or poor individual drug responses. This suggests that it is warranted to explore other mutations than those documented in animal worms.

## Introduction

The two benzimidazole (BZ) drugs, albendazole (ALB) and mebendazole (MEB), are the drugs of choice in school-based large-scale deworming programs against soil-transmitted helminths (STHs; which include the roundworms *Ascaris lumbricoides* (representative of Clade III), *Trichuris trichiura* (representative of Clade I) and the two hookworms, *Necator americanus* and *Ancylostoma duodenale* (representatives of Clade V)) [1,2]. During 2022 only, a BZ tablet was administered to 390 million school-aged children (SAC), covering 61.8% of the children at risk for the morbidity attributable to STH infections [3]. With the continued donation of both drugs by GlaxoSmithKline (ALB) and Johnson & Johnson (MEB), the World Health Organization (WHO) aims to reduce the prevalence of moderate-to-heavy intensity infections below 2% by 2030 [4]. However, the prospects of increased and continued drug pressure for a long period of time makes these public health programs highly vulnerable for the emergence of BZ resistance [5,6]. This is because ALB and MEB drugs share their mode of action (inhibition of the polymerisation of tubulin to microtubules by binding to the ß-tubulin) and when they are administered as a single oral fixed dose (ALB: 400 mg; MEB: 500 mg) [7,8], which is not the most optimal regimen for each of the different STH species [9–11].

BZ resistance is generally associated with single nucleotide polymorphisms (SNPs) in the gene encoding for ß-tubulin isotype 1 in Clade V roundworms in codons 167 (TTC to TAC), 198 (GAA to GCA) and/or 200 (TTC to TAC) [12,13]. In contrast to BZ resistance in animal STHs, which predominantly belong Clade V (e.g. trichostrongylids), and for which there is compelling evidence for the association between these SNPs and reduced therapeutic efficacy [12,13], little is known about these SNP occurrences in human STHs, and particularly in worms that are not Clade V (*Ascaris lumbricoides* and *Trichuris trichiura*), and their association with reduced efficacy of BZ drugs. Generally, this lack of evidence concerning human STHs, is multifactorial. First, studies reporting reduced therapeutic efficacy (commonly measured as reduction in egg excretion following deworming, ERR) as defined by WHO [14] are few in number (ALB: ERR*_Ascaris_* <85% [15, 16]; ERR_hookworms_ <80% [17]; ERR*_Trichiura_* <40% [10,16,18–21]; MEB: ERR_hookworms_ <60% [17,22]), sometimes flawed in study design (follow-up period too short to accurately assess drug efficacy [15,16,23]) and rarely followed by an assessment of SNPs [15,16].

Second, the few studies assessing SNPs were based on a small number of individuals per study area (≤ 10, [24–28]) or geographically confined to areas in STH endemic countries in Africa (Ghana [29], Ethiopia [30], Kenya [16,31], Rwanda [15], Tanzania [31–33] and Uganda [27]), Americas (Brazil, [34], Guatemala [35], Haiti [31,33,36], Honduras [24], Jamaica [25], Panama [16,36], Saint Lucia [25]) and Asia (China [25,27], Indonesia [37], Myanmar [38], Siri-Lanka [39], Thailand [25] and Vietnam [25]). Third, SNPs data were often collected applying a variety of study methodologies, impeding a ready interpretation and comparison of the study results. For example, different sample types (adult worms [24–28,30,32], larval stages [32,39] and pools of eggs [16,29,31,32–34,36,38,39]) originating from a wide spectrum of the population (children [15,24,26,27,30,31] or all ages [16,31,33,36,38]) that were examined by different molecular tools to assess SNPs (PCR-sequencing [15,24,25,31,37], PCR-RFLP [34], qPCR [32,29,38], pyrosequencing [16,31,33], isothermal nucleic acid amplification assays [36,39] and deep amplicon Illumina sequencing [30]). Too often the therapeutic drug efficacy is either not reported or determined by different egg counting methods [16], which further complicates comparison of the study results. Fourth, it has been shown that the essential ß-tubulin gene family is diverse between animal and human helminth species and between roundworms of different clades [1,30], and that not all ß-tubulin genes are associated with BZ resistance [30]. Finally, genes not encoding for tubulin may play a role in BZ resistance [40]. As a consequence, it remains unclear which genes to prioritize in future routine surveillance for emergence of BZ resistances in human STHs.

The overall aim of this study was to gain insights into the role of SNPs in the ß-tubulin gene, which cause BZ resistance in veterinary roundworms, as markers for BZ resistance in human STHs. For this, we first determined the analytical performance of previously described pyrosequencing assays for *A. lumbricoides*, *T. trichiura* and *N. americanus*, to detect and quantify SNPs of the ß-tubulin gene that appeared to most closely match the ß-tubulin isotype 1 gene of the veterinary roundworm *Haemonchus contortus*. Subsequently, we applied the assay on stool samples that were previously collected during three drug efficacy trials in Ethiopia, Lao PDR and Pemba Island (Tanzania). These trials were standardized in terms of egg count methods (Kato-Katz thick smear, Mini-FLOTAC, FECPAK^G2^ and qPCR), follow-up period (2 – 3 weeks), study population (SAC), the administered drug (single dose 400 mg ALB originating from the same manufacturer and batch) and statistical data analysis (group-based egg reduction rates based on arithmetic means). In addition, they were performed in areas exposed to different histories of BZ drug treatment and showed that efficacies were lowest at sites with a longer history of drug donation programs [18].

## Methods

### Ethics statement

The data were collected during clinical trials in Ethiopia, Lao PDR and Tanzania [18]. The study protocol for these trials was reviewed and approved by the institutional review board of the Faculty of Medicine and Health Sciences, and the University Hospital of Ghent University, Belgium (Ref. No B670201627755; 2016/0266) and by the responsible ethical committees associated with each trial site (Ethical Review Board of Jimma University, Jimma, Ethiopia: RPGC/547/2016; National Ethics Committee for Health Research (NECHR), Vientiane, Lao PDR: 018/NECHR; Zanzibar Medical Research and Ethics Committee, United Republic of Tanzania: ZAMREC/0002/February/2015). The trial was retrospectively registered on Clinicaltrials.gov (ID: NCT03465488) on March 7, 2018.

### Analytical performance of the pyrosequencing assays

#### PCR and pyrosequencing assays

We adapted previously described PCR and pyrosequencing assays [31,33] targeting β-tubulin genes identified in these previous studies to be homologs of Hco-tbb-iso-1. Briefly, primers for PCR and pyrosequencing (**Table 1**) were designed by Pyromark Q48 software. The PCR master mix contained per reaction 5 μL 10 x PCR buffer, 2 μL MgSO_4_ (50 mM), 1 μL dNTP (10 mM), 1 μL of bovine serum albumin (Sigma), 2 μL of each forward and reverse (biotinylated) primer (10 μM), 3 U Platinum High Fidelity Taq DNA polymerase (Invitrogen), and 4 μL sample DNA and distilled H_2_O to reach a final volume of 50 μL. Negative and positive controls (synthetic DNA sequences prepared as positive controls for the SNPs being investigated) were also included for quality control. The PCR reaction conditions were 94°C for 4 min, followed by 40 cycles at 94°C for 45 sec, 57-59 °C (see **Table 1**) for 45 sec and 68°C for 1 min and a final extension at 68°C for 10 min. PCR products were visualized after 2% acrylamide gel electrophoresis using ethidium bromide staining. Samples with the expected fragment size were subjected to pyrosequencing analysis. *Ascaris*, at position 167 was not analysed in this study as a previous study showed the presence of TAC codon at 167 had no relation to BZ resistance [16].

**Table 1.**
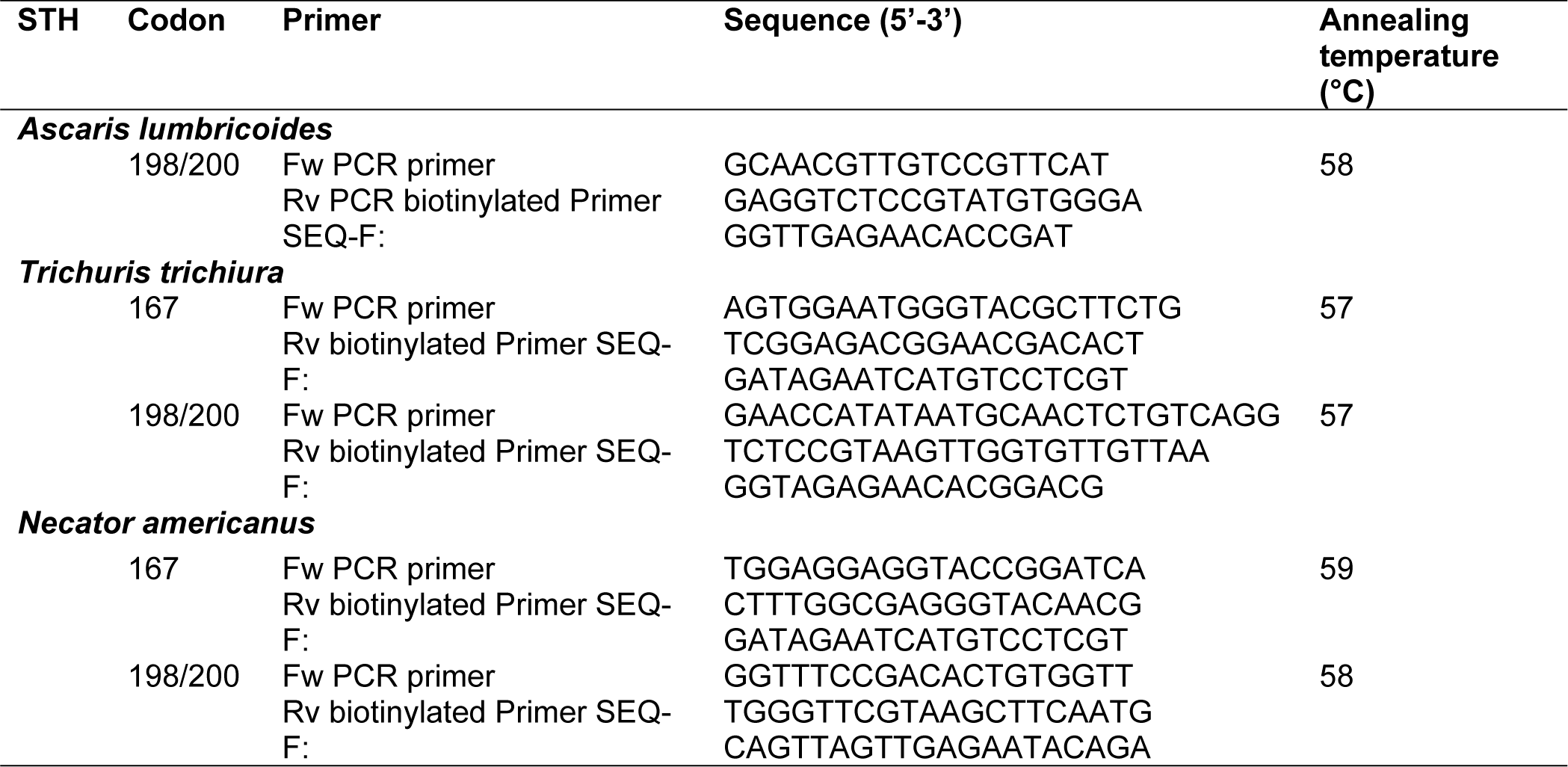
The soil-transmitted helminth (STH) specific primer sequences used in the PCR and pyrosequencing. *A. lumbricoides* (GenBank accession no. FJ501301.1; *Alu-bt-A* [30]), *T. trichiura* (GenBank accession no. AF034219.1), *N*. *americanus* (GenBank accession no. EF392851). Fw: forward; Rv: reverse, SEQ: sequencing.

#### Analytical performance parameters

For the pyrosequencing assays, we assessed four important analytical performance parameters, including the limit of blank (LoB), the limit of detection (LoD), and both the precision and the accuracy of the assay to quantify the proportion of mutant SNPs. To this end, we determined the proportion of both wildtype (WT) and mutant (MT) SNPs in samples containing a known ratio of MT to WT SNPs. For each codon, plasmids with the WT (codon 167: TTC; codon 198: GAA; codon 200: TTC) and MT sequence (codon 167: TAC; codon 198: GCA; codon 200: TAC)) were constructed (16 plasmids: 4 sequences for *A. lumbricoides* covering WT and MT SNPs at codons 198 and 200, and 12 sequences for *T. trichiura* and *N. americanus*, sequences covering the WT and MT SNPs at codons 167, 198 and 200). MT plasmids were engineered by site-directed mutagenesis. Primers for WT and MT plasmids (outer primers and inner primers carrying the mutant alleles) are shown in **Table 2**. Amplified WT or MT fragments were cloned into TOPO-TA-Cloning vector (Invitrogen). Plasmids were extracted and purified using the QIAprep spin miniprep kit (Qiagen) and subsequently sequenced by Sanger sequencing at the McGill University/Genome Quebec Innovation Centre, Montreal, Quebec (Canada). The quantity and purity of DNA was measured using a NanoDrop Spectrophotometer, ND-1000 (Implen, Munich, Germany). Diluted WT and MT plasmids were used for assay optimization and development, and to determine the LoD of the pyrosequencing assay.

**Table 2.**
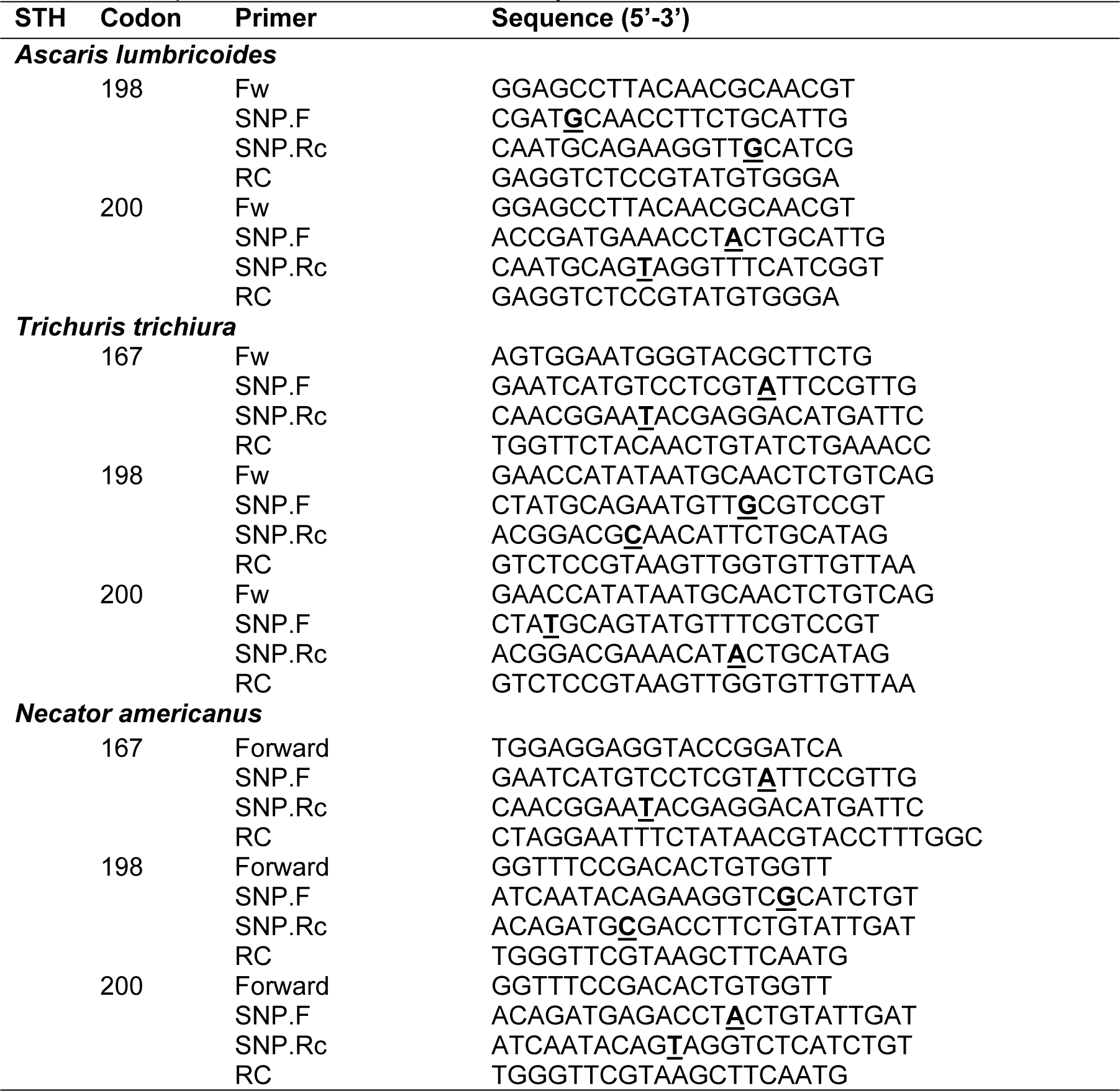
The soil-transmitted helminth (STH) specific primers for the wild and mutant plasmid constructs. The position of the SNP is indicated by a bold, underlined letter.

We tested 14 different ratios of MT to WT SNPs (MT:WT; 0:100, 1:99, 2:98, 3:97, 4:96, 5:95, 6:94, 7:93, 8:92, 9:91, 10:90; 20:80, 40:100, 100:0). All ratios were tested in triplicate, except for the 0:100 (MT:WT) where ten replicates were included. The LoB was determined for each SNP separately by the mean frequency of WT SNPs + 3 x standard deviation (SD) across the 0:100 replicates (n = 10). For the LoD, we first calculated the mean for each non-zero ratio of WT SNPs, and the median SD across all non-zero SD. Subsequently, we constructed a normal distribution with μ (= mean) and σ (= median SD) and determined the probability (= analytical sensitivity) of observing a frequency of WT SNP higher than the LoB. The LoD equalled to the lowest ratio of WT SNPs for which the analytical sensitivity was ≥95%.

For the quantitative test results, we first verified whether the observed proportions of MT SNPs were biased (measure of accuracy). In case of different frequencies (bias), we fitted linear regression models for each SNP with the observed frequency of MT SNPs as the outcome and the true underlying ratio of each of the MT SNPs as predictive variable. The precision of the pyrosequencing assay was estimated by the coefficient of variance.

### Occurrence of SNPs in ß-tubulin in stool samples

#### Origin of the field samples

The stool samples were collected as part of three clinical trials that were designed to assess therapeutic efficacy of a single oral dose of 400 mg ALB against STH infections in SAC measured by a variety of diagnostic methods (Kato-Katz, Mini-FLOTAC, FECPAK^G2^ and qPCR) at Lao PDR, Ethiopia and Pemba Island (Tanzania). Vlaminck et al. previously provided a detailed description of the study design [18, 41]. In brief, schools were visited by the local principal investigator and a team of field officers, who explained the planned trial and sampling method to the parents, teachers and the children. At baseline, demographic data of the SAC whom parents consented to allow their children to join the trial were collected, and they were asked to provide a fresh stool sample. Upon delivery of the stool samples, they were treated with a single oral dose of 400 mg of ALB under supervision. Fresh samples were processed the same day on site by duplicate Kato-Katz, Mini-FLOTAC and FECPAK^G2^. For qPCR, an aliquot of the samples was preserved in ethanol (0.5 g of stool in 1 mL absolute ethanol), which were subsequently shipped to the Laboratory of Parasitology (Belgium) and the Elisabeth-Tweesteden Hospital (Netherlands) for both DNA extraction and qPCR analysis [42]. Fourteen to 21 days after drug administration, a second stool sample was collected from all the children that were found positive for any STH at baseline. Stool samples collected at follow-up were again processed with the previously described methods. The details on the demographics, the STH status and the group-based therapeutic drug efficacy across the study sites results are described by Vlaminck et al. 2019 [18]. The analysis of the individual responses to the treatment is reported by Walker et al., 2022 [43].

#### Selection of the samples

Two different sample sets were created. One set of samples was used to address the question whether there is an association between frequency of mutant SNPs and history of drug donation programs (drug history sample set), while another set was used to verify whether poor drug responses is associated with a higher frequency of mutant SNPs (drug response data set). The drug history sample set only includes baseline samples, while the drug response sample set includes both baseline and follow-up samples. In order to increase the probability of sufficient target DNA prior to PCR and pyrosequencing, we only considered subjects for which the mean fecal egg counts (FECs) across the different egg counting methods (single and duplicate Kato-Katz thick smear, Mini-FLOTAC and FECPAK^G2^) at baseline were ≥150 eggs per gram of stool (EPG). For the follow-up samples, only samples in which at least one egg was detected by one of the four microscopic methods were investigated.

For the drug history sample set, it was envisioned to analyse at least 60 baseline samples per STH species and study site, if possible, resulting in 360 analyses for *Ascaris* (2 codons x 3 study sites x 60 samples) and 540 for *Trichuris* and *Necator* (3 codons x 3 study sites x 60 sample). As multiple schools were involved at the different study sites (Ethiopia: 2, Lao PDR: 5 and Pemba Island (Tanzania): 4), we ensured that the sample set represented each school and that the number of samples included was proportionally to number of samples collected at the different schools. In other words, proportionally more samples were selected from schools that delivered more cases.

For the drug response sample set, it was envisioned to have at least 60 subjects per STH species and study site, of whom ≥30 subjects were cured following drug administration, ≥10 showed a satisfactory drug response (good responders), ≥10 a doubtful drug response (poor responders) and ≥10 a reduced efficacy (non-responders). The assessment of the individual drug response was based on the individual reduction in egg counts following drug administration, where the egg counts equalled to mean FECs (in EPG) across the different microscopic methods. To classify the good, poor and non-responders, we applied the WHO criteria to classify the efficacy of single oral dose 400 mg ALB (**Table 3**) on the individual ERRs [14].

**Table 3.** Classification of the efficacy of a single oral dose 400 mg albendazole against soil-transmitted helminths. The efficacy was measured as the reduction in mean fecal egg counts following drug administration (ERR). These thresholds were recommended by the World Health Organization for a single Kato-Katz.

| STH | Satisfactory<br>(Good responders) | Doubtful<br>(Poor responders) | Reduced<br>(Non-responders) |
| --- | --- | --- | --- |
| *Ascaris lumbricoides* | $\geq 95\%$ | $85\% \leq \text{ERR} < 95\%$ | $< 85\%$ |
| *Trichuris trichiura* | $\geq 50\%$ | $40\% \leq \text{ERR} < 50\%$ | $< 40\%$ |

| Hookworm | ≥90% | 80%≤ ERR <90% | <80% |
| --- | --- | --- | --- |

To minimize the required resources to obtain these numbers of cases for each of the four classes of drug response (cured subjects, good, poor and non-responders), we already considered the subjects selected for the drug history sample set. Subsequently, subjects were added until the required number of samples was obtained. Similarly, as for the drug history sample set, we ensured that the sample set represented each school and that the number of samples included was proportionally to number at the different schools. In other words, proportionally more samples were selected from schools that delivered more cases. Note that we did not conduct any sample size calculation to test of the aforementioned hypothesis.

#### Sample processing

In brief, DNA from all samples were first subjected to the PCR protocols (*Ascaris*: 1, *Trichuris*: 2 and *Necator*: 1). In case of insufficient DNA amplification (absence or weak band on gel), we performed a nested PCR, using 1 μL of the initial PCR product as DNA template and the same primers. Only PCR products with sufficient DNA amplification (strong bands on gel) were further analysed with the corresponding pyrosequencing assays. To also verify the repeatability of pyrosequencing test results on stool samples, we subjected the PCR products of the first 50 *Trichuris* and *Necator* samples twice to the pyrosequencing assays. The repeatability of the pyrosequencing assays for *Ascaris* was not assessed. This is because we prioritized the analysis of *Trichuris* and *Necator* samples (for these STHs the therapeutic efficacy was lowest at sites with a longer history of drug donation programs) and because the analysis of the PCR products in duplicates for the other STHs indicated that pyrosequencing results were highly repeatable. All the frequencies of MT SNPs obtained following pyrosequencing were corrected to account for an overestimation, applying the formulae described in **Table 4** to gain more accurate estimates of the true underlying frequency (results indicated an overestimation of the frequency of MT SNPs).

**Table 4.** The formulae used to provide more accurate estimates of true underlying frequency of mutant SNPs. For each codon and STH species, two formulae were used. The first accounts for the limit of the blank, the corrected frequency of mutant SNP (*y_cor_*) being set at zero when the observed frequency (*y_obs_*) did not exceed the limit of the blank. In all other cases, the *y_cor_* was determined as a function of *y_obs_*. For this, the outcome of the linear regression model (*y_obs_ = a + b*(true frequency of SNPs*) was converted into true underlying frequency of SNPs *=* (*y_obs_ – a*)*/b* ≈ *y_cor_*).

| STH | Codon | Formula for $y_{cor}$ based on $y_{obs}$ |
| --- | --- | --- |
| *Ascaris* | 198 | If $y_{obs} \leq 3.08$ then $y_{cor} = 0$ |
| | | if $y_{obs} > 3.08$ then $y_{cor} = (y_{obs} - 1.10)/0.97$ |
| | 200 | If $y_{obs} \leq 2.95$ then $y_{cor} = 0$ |
| | | if $y_{obs} > 2.95$ then $y_{cor} = (y_{obs} - 1.12)/0.97$ |
| *Trichuris* | 167 | If $y_{obs} \leq 3.06$ then $y_{cor} = 0$ |
| | | if $y_{obs} > 3.06$ then $y_{cor} = (y_{obs} - 0.66)/0.97$ |
| | 198 | If $y_{obs} \leq 3.46$ then $y_{cor} = 0$ |
| | | if $y_{obs} > 3.46$ then $y_{cor} = (y_{obs} - 1.27)/0.93$ |
| | 200 | If $y_{obs} \leq 2.85$ then $y_{cor} = 0$ |
| | | if $y_{obs} > 2.85$ then $y_{cor} = (y_{obs} - 1.57)/0.98$ |
| *Necator* | 167 | If $y_{obs} \leq 5.71$ then $y_{cor} = 0$ |
| | | if $y_{obs} > 5.71$ then $y_{cor} = (y_{obs} - 0.57)/0.98$ |
| | 198 | If $y_{obs} \leq 3.41$ then $y_{cor} = 0$ |
| | | if $y_{obs} > 3.41$ then $y_{cor} = (y_{obs} - 1.83)/0.96$ |
| | 200 | If $y_{obs} \leq 3.06$ then $y_{cor} = 0$ |
| | | if $y_{obs} > 3.06$ then $y_{cor} = (y_{obs} - 1.99)/0.97$ |

#### Statistical data analysis

##### Drug history sample set

We verified any association between both the occurrence (presence/absence of MT SNPs within a sample) and the frequency of MT SNPs (the proportion of MT SNPs within a sample), and history of drug donation programs. To this end, we explored differences in both the occurrence of MT SNPs (Fisher’s exact test) and frequency of MT SNPs (the Kruskal-Wallis) within a sample across the three study sites (Ethiopia: short history of drug donation programs (since 2014); Lao PDR: moderate history of drug donation program (since 2005); Pemba Island: long history of drug donation program (since 1999)) for each codon and STH species In case of significant difference (*p* <0.05), we continued with pair-wise comparisons between study sites applying Fisher’s exact test (occurrence of MT SNPs) and the Mann-Whitney-U (frequency of MT SNPs).

##### Drug response sample set

For the association between MT SNPs and the individual response we explored the difference in both the occurrence of MT SNPs (Fisher’s exact test) and frequency of MT SNPs (the Kruskal-Wallis) within a sample across the different levels of individual treatment responses (cured subjects, good, poor and non-responders) for each codon and STH species, respectively. We also explored whether there was an increase in frequency in MT SNPs following treatment. For this we applied a paired Kruskal-Wallis test to detect any difference in the frequency in MT SNPs in baseline and follow-up samples of the same individuals.

##### Repeatability of pyrosequencing test results on field samples

Bland-Altman plots were made for each of the assays. In these plots, the difference in frequency of MT SNPs in the two replicate DNA concentrations was plotted as a function of the mean frequency of mutant SNPs. In addition, the limit of agreement (range of the difference in frequency of mutant SNPs between replicates that cover 95% of the values) and their corresponding 95% confidence intervals (95% CI) were determined by bootstrap analysis (5,000 iterations).

## Results

### Analytical performance of the pyrosequencing assays

Overall, the LoB was ∼3.5% for all codons and STHs, except for *Necator*. For this STH, a LoB of 5.7% was observed for codon 167. Generally, the LoD increased as a function of LoB, ranging from 2% to 7%. For most codons, the LoD was 2% to 4%, whereas it was 7% for codon 167 of *Necator*. As shown in the **Figs 1** to **3**, the pyrosequencing overestimated the true underlying frequency (the lower limit of the 99.7% confidence interval did not include the line of equivalence in several cases). Based on the LoB and the outcome of the linear regression models we used the formula described in **Table 4** to correct the observed frequencies to more accurately estimate the true underlying frequencies of MT SNPs. Generally, the precision (coefficient of variance) of the assays decreased as a function of increasing frequency of true underlying frequency of mutant SNPs. Where the coefficient of variance exceeded 20% when the frequency of SNPs was below 3% for *Ascaris* and *Necator*, and 4% for *Trichuris*, whereas a coefficient of variance of less than 10% across all codons was observed once the frequency was at least 9% (*Trichuris*), 10% (*Ascaris*) and 20% (*Necator*; **Table S1**).

**Fig 1.**
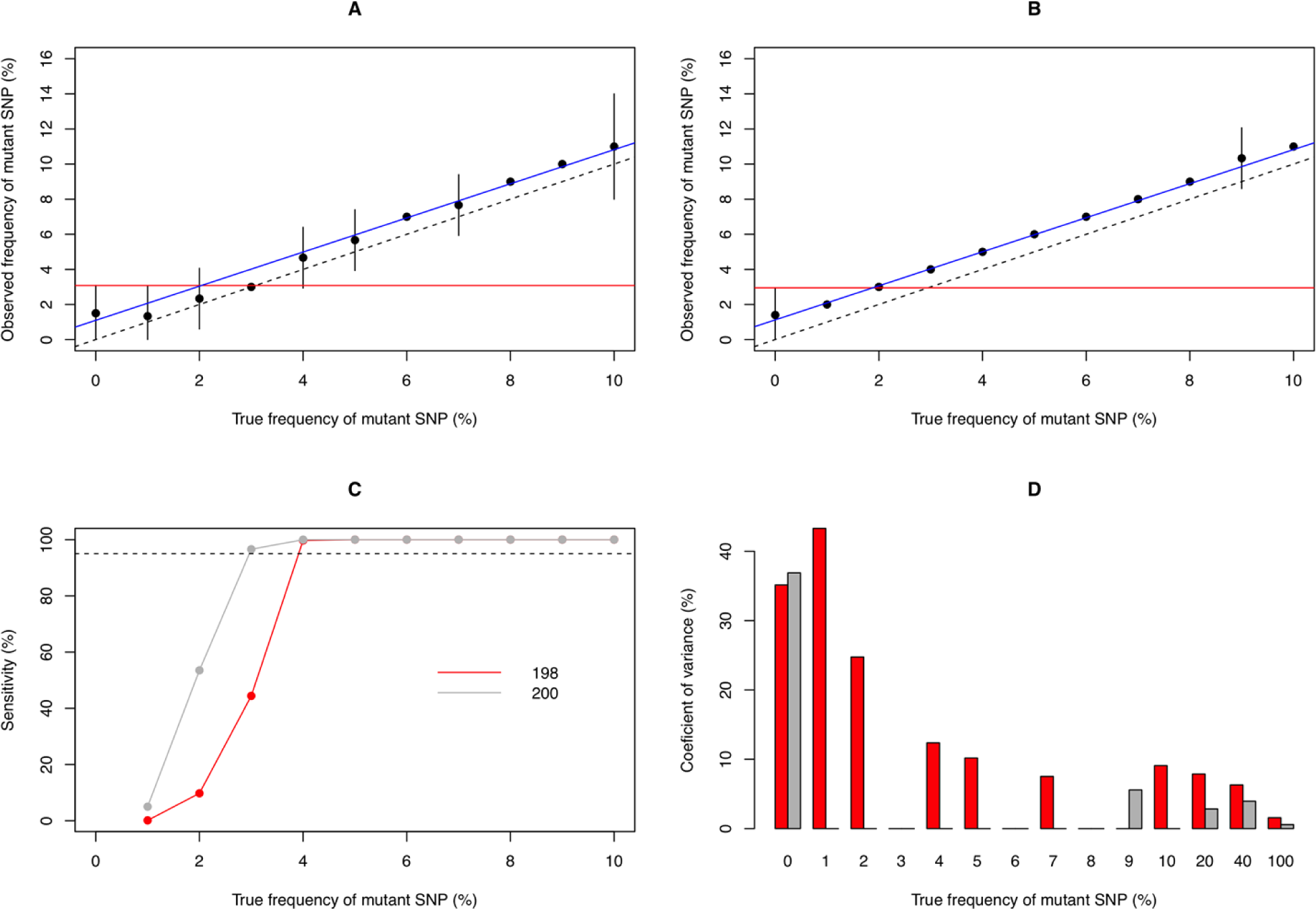
The analytical performance of the pyrosequencing assay for SNPs in the ß-tubulin gene of *Ascaris*. The scatter plots in **Panels A** and **B** illustrate the observed frequency of mutant SNPs for 11 true frequencies of mutant (0 – 10%) for codon 198 (**Panel A**) and codon 200 (**Panel B**). In these plots the dots represent the mean observed frequency of mutant SNPs for a given true underlying frequency, the whiskers represent 99.7% of the potential values (mean +/− 3 x standard deviation) and the limit of blank is represented by a red line. The dashed line represents the line of equivalence, whereas the blue line represents the outcome of the linear model with the observed frequency as outcome variable and the true underlying frequency as predictive variable. **Panel C** illustrates the analytical sensitivity of the assay for the detection of mutant SNPs, whereas **Panel D** illustrates the coefficient of variance (red: codon 198, grey: codon 200).

**Fig 2.**
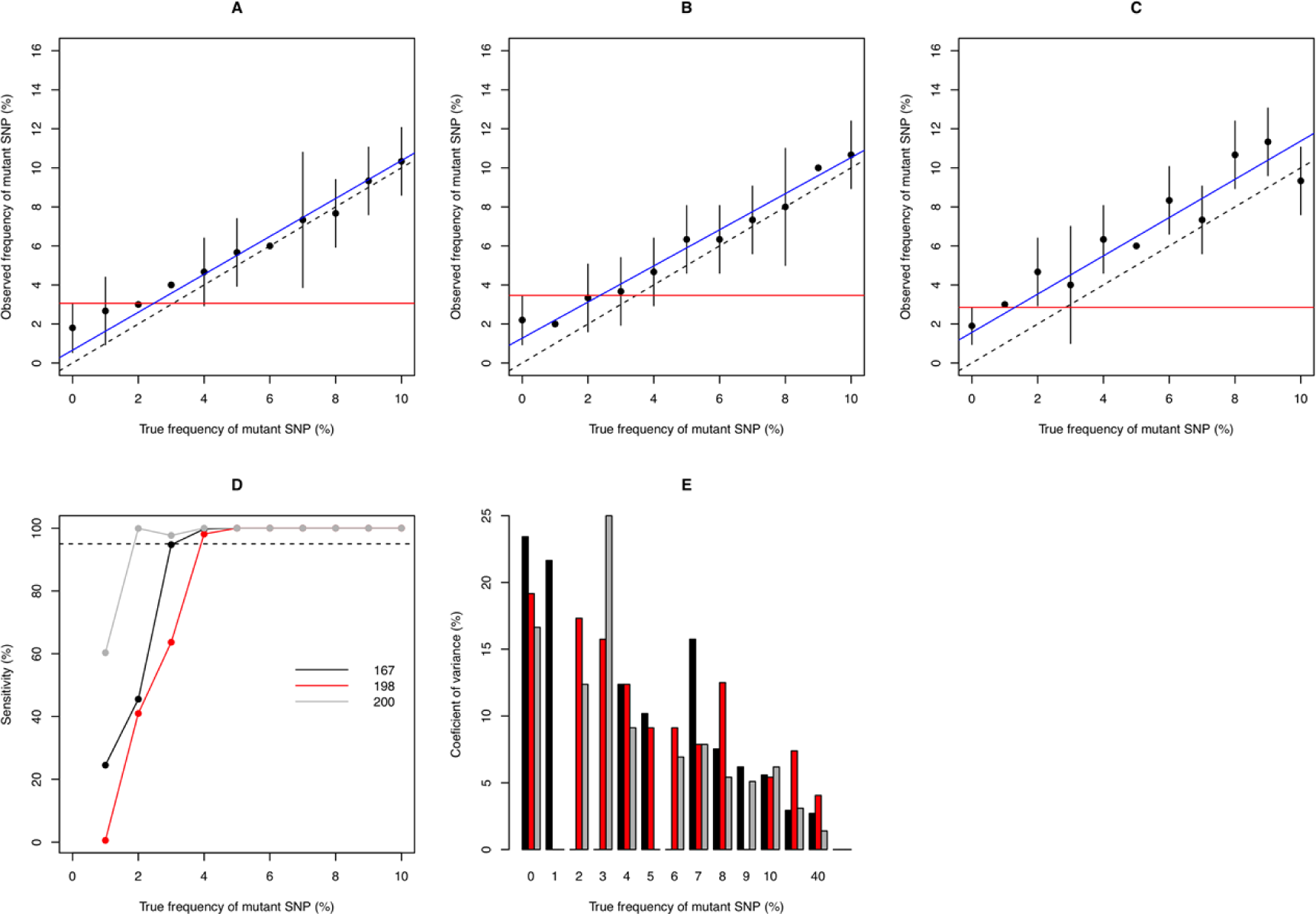
The analytical performance of the pyrosequencing assay for SNPs in the ß-tubulin gene of *Trichuris*. The scatter plots in **Panels A** to **C** illustrate the observed frequency of mutant SNPs for 11 true frequencies of mutant (0 – 10%) for codon 167 (**Panel A**), codon 198 (**Panel B**) and codon 200 (**Panel C**). In these plots the dots represent the mean observed frequency of mutant SNPs for a given true underlying frequency, the whiskers represent 99.7% of the potential values (mean +/− 3*standard deviation) and the limit of blank is represented by a red line. The dashed line represents the line of equivalence, whereas the blue line represents the outcome of linear model with the observed frequency as outcome variable and the true underlying frequency as predictive variable. **Panel D** illustrates the analytical sensitivity of the assay for the detection of mutant SNPs, where **Panel E** illustrates the coefficient of variance (black: codon 167, red: codon 198, grey: codon 200).

**Fig 3.**
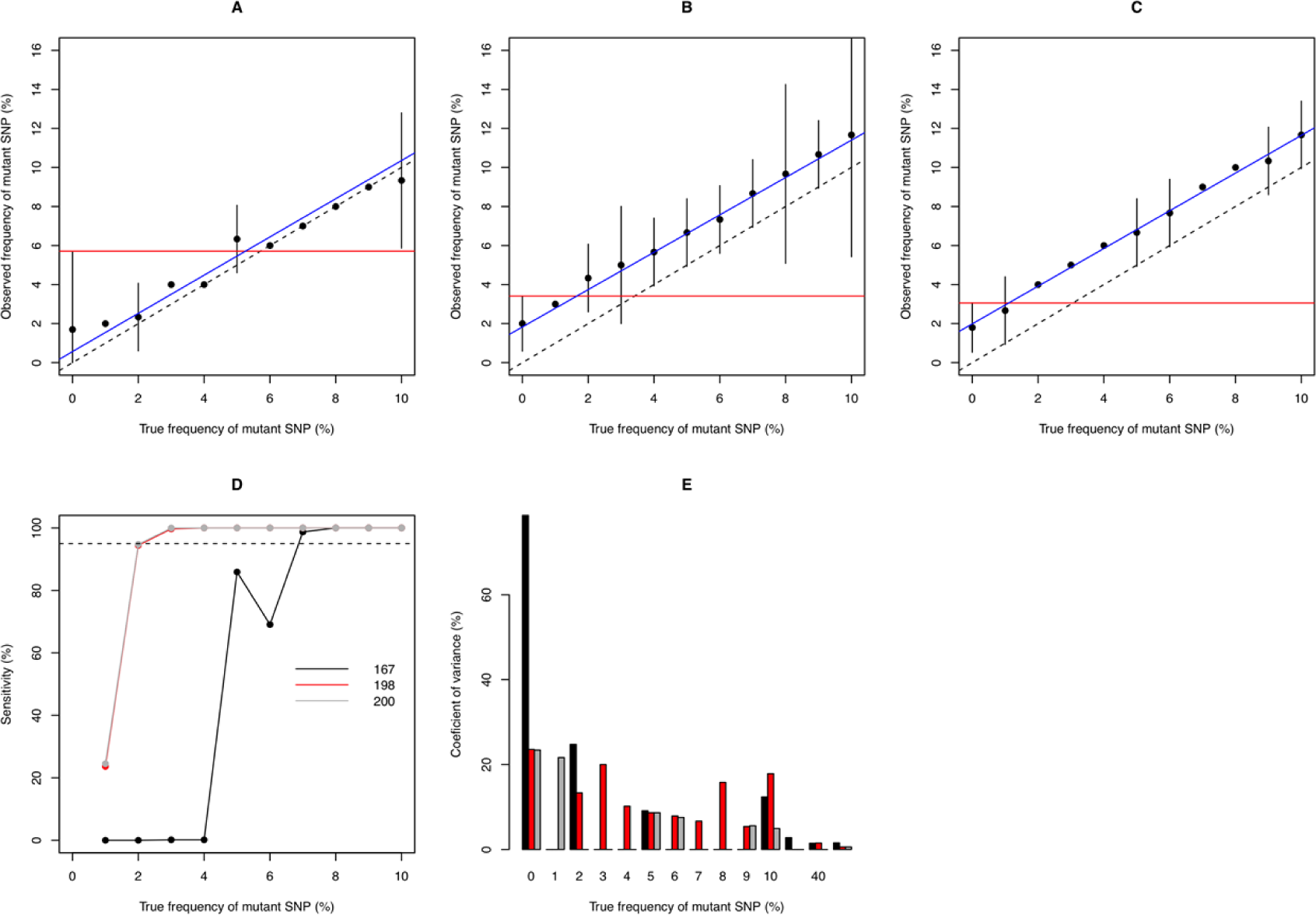
The analytical performance of the pyrosequencing assay for SNPs in the ß-tubulin gene of *Necator*. The scatter plots in **Panels A** to **C** illustrate the observed frequency of mutant SNPs for 11 true frequencies of mutant (0 – 10%) for codon 167 (**Panel A**), codon 198 (**Panel B**) and codon 200 (**Panel C**). In these plots the dots represent the mean observed frequency of mutant SNPs for a given true underlying frequency, the whiskers represent 99.7% of the potential values (mean +/− 3*standard deviation) and the limit of blank is represented by a red line. The dashed line represents the line of equivalence, whereas the blue line represents the outcome of linear model with the observed frequency as outcome variable and the true underlying frequency as predictive variable. **Panel D** illustrates the analytical sensitivity of the assay for the detection of mutant SNPs, where **Panel E** illustrates the coefficient of variance (black: codon 167, red: codon 198, grey: codon 200).

### Occurrence of SNPs in ß-tubulin associated with BZ resistance in stool samples

#### Association with history of drug donation programs

A total of 827 baseline samples were available from an equal number of subjects, resulting in 436 *Ascaris* (Ethiopia: 141, Lao PDR: 108, Pemba Island: 187), 541 *Trichuris* (Ethiopia: 153, Lao PDR: 109, Pemba Island: 279), and 500 hookworm cases (Ethiopia: 111, Lao PDR: 268, Pemba Island: 121). The majority of these subjects (645; 78.0%) excreted on average ≥ 150 EPG. Yet, it became clear that the target of 60 subjects for each study site per STH was difficult to meet. In total 156 *Ascaris* (Ethiopia: 58, Laos PDR: 59, Pemba Island: 49), 119 *Trichuris* (Ethiopia: 31, Laos PDR: 34, Pemba Island: 54) and 145 *Necator* DNA samples (Ethiopia: 36, Laos PDR: 58, Pemba Island: 51) were processed with PCR. For all STHs, ∼75% of the DNA samples subjected to PCR resulted in visible amplicons for at least one of the pyrosequencing assays (*Ascaris*: 76.6%, *Trichuris*: 73.1% and *Necator*: 79.3%). In all other cases, no visible amplicon was obtained.

**Table 5** further details the occurrence of MT SNPs across the different study sites with varying history of drug donation programs (Ethiopia, Lao PDR, Pemba Island: respectively short, moderate and long history of drug donation) for each codon and STH species separately. While MT SNPs in codon 200 were detected in half of the analysed *Trichuris* samples (33 out of 65), the proportion of the analysed samples containing MT SNPs did not exceed 15% for all other codons and STH species. Generally, the frequency of MT SNPs within a sample was low. The highest ratio was observed in a *Trichuris* sample for which the adjustedMT SNP frequency at codon 167 equalled 26.1%. In the remaining samples the frequency of MT SNPs did not exceed 15% (2.1% - 12.7%).

**Table 5.** The occurrence and ratio of mutant SNPs across three codons at the three study sites with varying history of drug pressure. N_PCR_: number of DNA samples subjected to PCR; N_PCR+_: number of DNA samples that tested positive PCR; N_MT+_: number of DNA samples in which mutant (MT) single nucleotide polymorphisms (SNPs) are detected; min – max adjusted frequency of MT SNPs: minimum and maximum frequency of MT SNPs. The frequencies are adjusted based on the formulae in Table 4; _ not available.

|  | Codon 167 |  |  |  | Codon 198 |  |  |  | Codon 200 |  |  |
| --- | --- | --- | --- | --- | --- | --- | --- | --- | --- | --- | --- |
|  | N<sub>PCR</sub> | N<sub>PCR+</sub> | N<sub>MT+</sub> | Min – max<br>adjusted<br>frequency of MT<br>SNPs | N<sub>PCR+</sub> | N<sub>MT+</sub> | Min – max<br>adjusted<br>frequency of<br>MT SNPs |  | N<sub>PCR+</sub> | N<sub>MT+</sub> | Min – max<br>adjusted<br>frequency of<br>MT SNPs |
| ***Ascaris*** |  |  |  |  |  |  |  |  |  |  |  |
| Ethiopia | 57 | – | – | – | 39 | 0 | – |  | 39 | 2 | 1.9 – 9.1 |
| Lao PDR | 54 | – | – | – | 39 | 1 | 18.5 |  | 39 | 4 | 1.9 – 11.2 |
| Pemba<br>Island | 45 | – | – | – | 40 | 0 | – |  | 40 | 2 | 1.9 – 4.0 |
| **Total** | **156** |  |  |  | **118**<br>**(75.6%)** | **1**<br>**(1.0%)** | **18.5** |  | **118**<br>**(75.6%)** | **8**<br>**(6.7%)** | **1.9 – 11.2** |
| ***Trichuris*** |  |  |  |  |  |  |  |  |  |  |  |
| Ethiopia | 31 | 8 | 0 | – | 5 | 1 | 2.9 |  | 5 | 5 | 1.5 – 9.6 |
| Lao PDR | 34 | 20 | 0 | – | 21 | 1 | 2.9 |  | 21 | 11 | 1.5 – 12.7 |
| Pemba<br>Island | 54 | 35 | 3 | 3.4 – 26.1 | 39 | 4 | 2.9 – 8.3 |  | 39 | 17 | 1.5 – 3.5 |
| **Total** | **119** | **63**<br>**(52.9%)** | **3**<br>**(4.8%)** | **3.4 – 26.1** | **65**<br>**(54.6%)** | **6**<br>**(9.2%)** | **2.9 – 8.3** |  | **65**<br>**(54.6%)** | **33**<br>**(50.8%)** | **1.5 – 12.7** |
| ***Necator*** |  |  |  |  |  |  |  |  |  |  |  |
| Ethiopia | 36 | 17 | 1 | 5.5 | 7 | 2 | 3.3 – 4.3 |  | 7 | 2 | 2.1 – 3.1 |
| Lao PDR | 58 | 50 | 0 | – | 48 | 3 | 2.3 – 4.3 |  | 48 | 5 | 2.1 – 3.1 |
| Pemba<br>Island | 51 | 39 | 1 | 11.7 | 43 | 9 | 2.3 – 5.4 |  | 43 | 2 | 2.1 – 3.1 |
| **Total** | **145** | **106**<br>**(73.1)** | **2**<br>**(1.4%)** | **5.5 – 11.7** | **98**<br>**(67.6%)** | **14**<br>**(14.2%)** | **2.3 – 5.4** |  | **98**<br>**(67.6%)** | **9**<br>**(9.2%)** | **2.1 – 3.1** |

The Fisher’s exact tests revealed a significant and a marginal difference in the occurrence of MT SNPs across the different study sites for *Necator* at codon 198 (Ethiopia: 2/7; Lao PDR: 3/48; Pemba Island: 9/43, *p* = 0.042) and codon 200 (Ethiopia: 2/7; Lao PDR: 5/48; Pemba Island: 2/43, *p* = 0.10), respectively. The pair-wise comparisons for codon 198 revealed a marginal difference between Lao PDR and Pemba Island (*p* = 0.061) only. Similarly, the Kruskal-Wallis tests revealed a significant and a marginal difference in the adjusted frequency of MT SNPs within samples across the different study sites for *Trichuris* at codon 200 (*p* <0.001) and for *Necator* at codon 198 (*p* = 0.065), respectively. The pair-wise comparison for codon 200 of *Trichuris* indicated that the frequency was significantly higher in Ethiopia (median adjusted ratio: 3.5%) compared to both Lao PDR (median adjusted frequency: 1.5%; *p* = 0.012) and Tanzania (median adjusted frequency: 0.0%; *p* = 0.002).

#### Association with therapeutic drug response

Here too it became clear that it was impossible to achieve the target sample size (≥30 cured subjects, ≥10 good responders, ≥10 poor responders and ≥10 non-responders) for each STH and study site separately, and that a PCR amplicon was not always visible for all codons in both samples for each subject. We therefore opted to reduce the number response levels from four to two by combining cured subjects and good responders into ‘satisfactory’ response, and the poor and non-responders into ‘poor-to-reduced’ response.

**Table 6** further details the occurrence of MT SNPs across the different treatment response levels for each codon and STH species separately, with the aim to assess whether the occurrence or the higher frequencies of MT SNPs at baseline is associated with more poor-to-reduced treatment responses. The output of the statistical analysis indicates no significant difference in both the occurrence (Fisher’s exact test) and the adjusted frequency of MT SNPs (Mann-Witney test) between subjects with a satisfactory and those with a poor-to-reduced treatment response.

**Table 6.** The occurrence and frequency of mutant SNPs across three codons at baseline between individuals with a varying response. N_PCR_: number of DNA samples subjected to PCR; N_PCR+_: number of DNA samples that tested positive PCR; N_MT+_: number of DNA samples in which mutant (MT) single nucleotide polymorphisms (SNPs) are detected; min – max adjusted frequency of MT SNPs: minimum and maximum frequency of MT SNPs. The frequencies are adjusted based on the formulae in Table 4; _ not available.

|  | Codon 167 |  |  |  | Codon 198 |  |  |  | Codon 200 |  |  |
| --- | --- | --- | --- | --- | --- | --- | --- | --- | --- | --- | --- |
|  | N<sub>PCR</sub> | N<sub>PCR+</sub> | N<sub>MT+</sub> | Min – max adjusted frequency of MT SNPs | N<sub>PCR+</sub> | N<sub>MT+</sub> | Min – max adjusted frequency of MT SNPs |  | N<sub>PCR+</sub> | N<sub>MT+</sub> | Min – max adjusted frequency of MT SNPs |
| ***Ascaris*** |  |  |  |  |  |  |  |  |  |  |  |
| Satisfactory response | 171 | – | – | – | 131 | 1 | 18.5 |  | 131 | 8 | 1.9 – 11.2 |
| Poor-to-reduced response | 6 | – | – | – | 6 | 0 | – |  | 6 | 0 | 0 |
| **Total** | **177** |  |  |  | **137 (77.4%)** | **1 (1.0%)** | **14.3 – 18.5** |  | **137 (77.4%)** | **8 (5.8%)** | **1.9 – 11.2** |
| ***Trichuris*** |  |  |  |  |  |  |  |  |  |  |  |
| Satisfactory response | 91 | 51 | 1 | 3.4 | 57 | 4 | 2.9 – 4.0 |  | 57 | 26 | 1.5 – 12.7 |
| Poor-to-reduced response | 55 | 31 | 2 | 5.5 – 26.1 | 32 | 4 | 2.9 – 8.3 |  | 32 | 14 | 1.5 – 3.5 |
| **Total** | **146** | **86 (58.9%)** | **3 (3.5%)** | **3.4 – 26.1** | **89 (60.9%)** | **8 (9.0%)** | **2.9 – 8.3** |  | **89 (60.9%)** | **40 (44.9%)** | **1.5 – 12.7** |
| ***Necator*** |  |  |  |  |  |  |  |  |  |  |  |
| Satisfactory response | 147 | 120 | 3 | 5.5 – 11.6 | 108 | 14 | 2.3 – 5.4 |  | 108 | 9 | 2.1 – 6.2 |
| Poor-to-reduced response | 39 | 31 | 0 | – | 34 | 4 | 2.3 – 4.3 |  | 34 | 1 | 3.1 |
| **Total** | **186** | **151 (81.2%)** | **3 (2.0%)** | **5.5 – 11.6** | **142 (76.3%)** | **18 (12.7%)** | **2.3 – 5.4** |  | **142 (76.3%)** | **10 (7.0%)** | **2.1 – 6.2** |

In **Fig 4** we explored whether the adjusted frequency of MT SNPs increased following drug administration for *Trichuris* and *Necator*. There was a significant increase in the adjusted ratio of MT SNP at codon 200 for *Trichuris* but not for *Necator*. There were no significant changes in MT frequencies at codons 167 and 198 for either of these species. For *Ascaris* there were too few subjects for which a complete analysis at both baseline and follow-up samples was available (when subjects were cured following treatment, no follow-up sample was preserved in the field).

**Figure 4.**
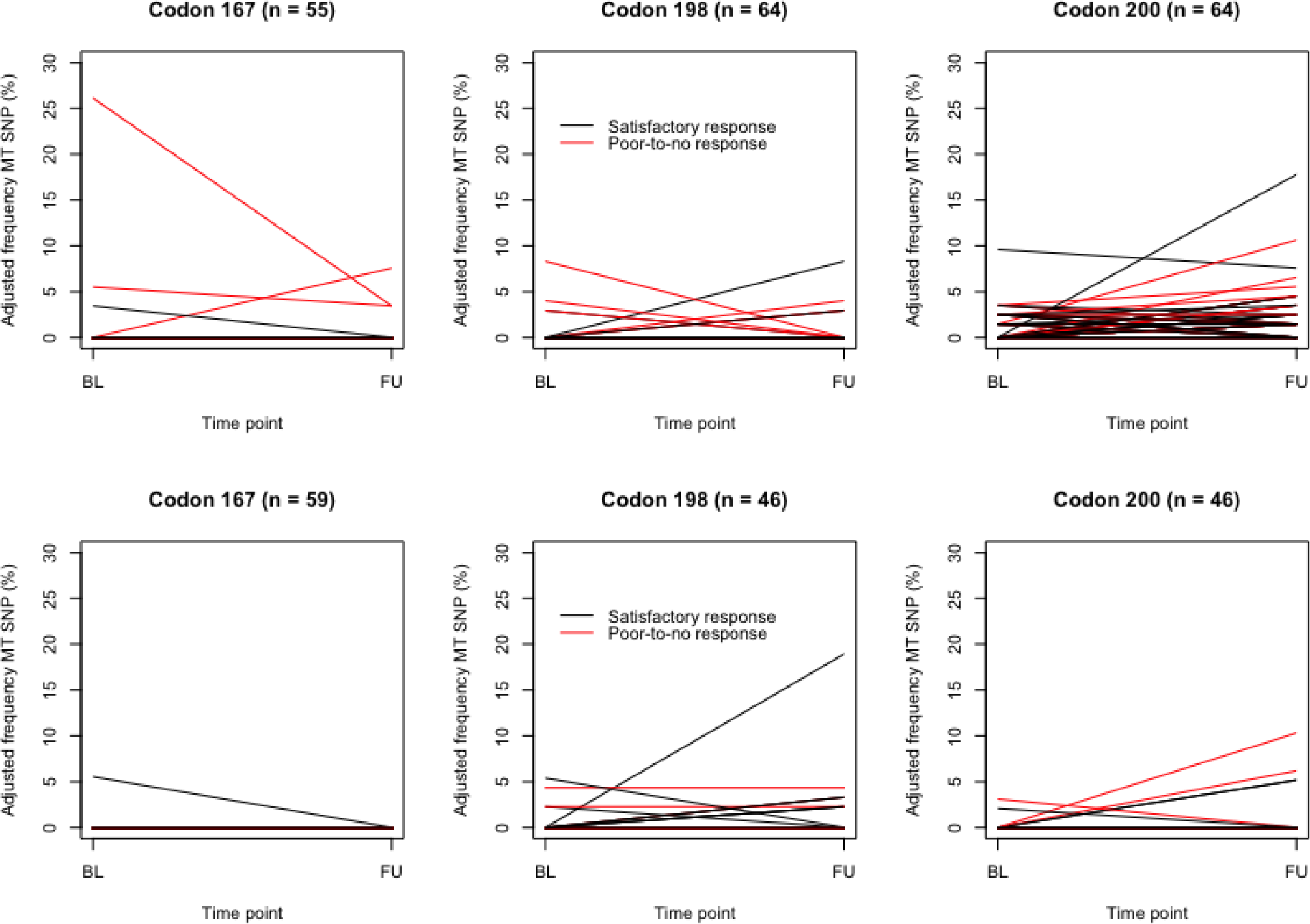
The change in adjusted frequency of mutant SNPs following a drug administration for *Trichuris* and *Necator*. This figure illustrates the change in adjusted mutant ratio (MT) SNPs in the ß-tubulin isotype 1 for *Trichuris* (**top row**) and *Necator* (**bottom row**) for codons 167, 198 and 200 before (BL) and after (FU) administration of a single oral dose of 400 mg albendazole. The black lines represent subjects with a satisfactory response, while the red lines represent those that poorly or did not respond to the treatment.

#### Repeatability of pyrosequencing assays on field samples

**Fig 5** displays the Bland-Altman plots based on the output of all pyrosequencing assays (no per codon analysis was presented) for both *Trichuris* and *Necator,* separately. These plots indicate that the difference in output between replicates is expected to fall between −2.1 (95%CI: −2.9; −1.5) and 3.0 (95%CI: 1.1; 3.4) for *Trichuris,* and between −2.1 (95%CI: −2.3; 0.0) and 3.2 (95%CI: 2.2; 3.3) for *Necator.* All the data is made available in **Info S1.**

**Figure 5.**
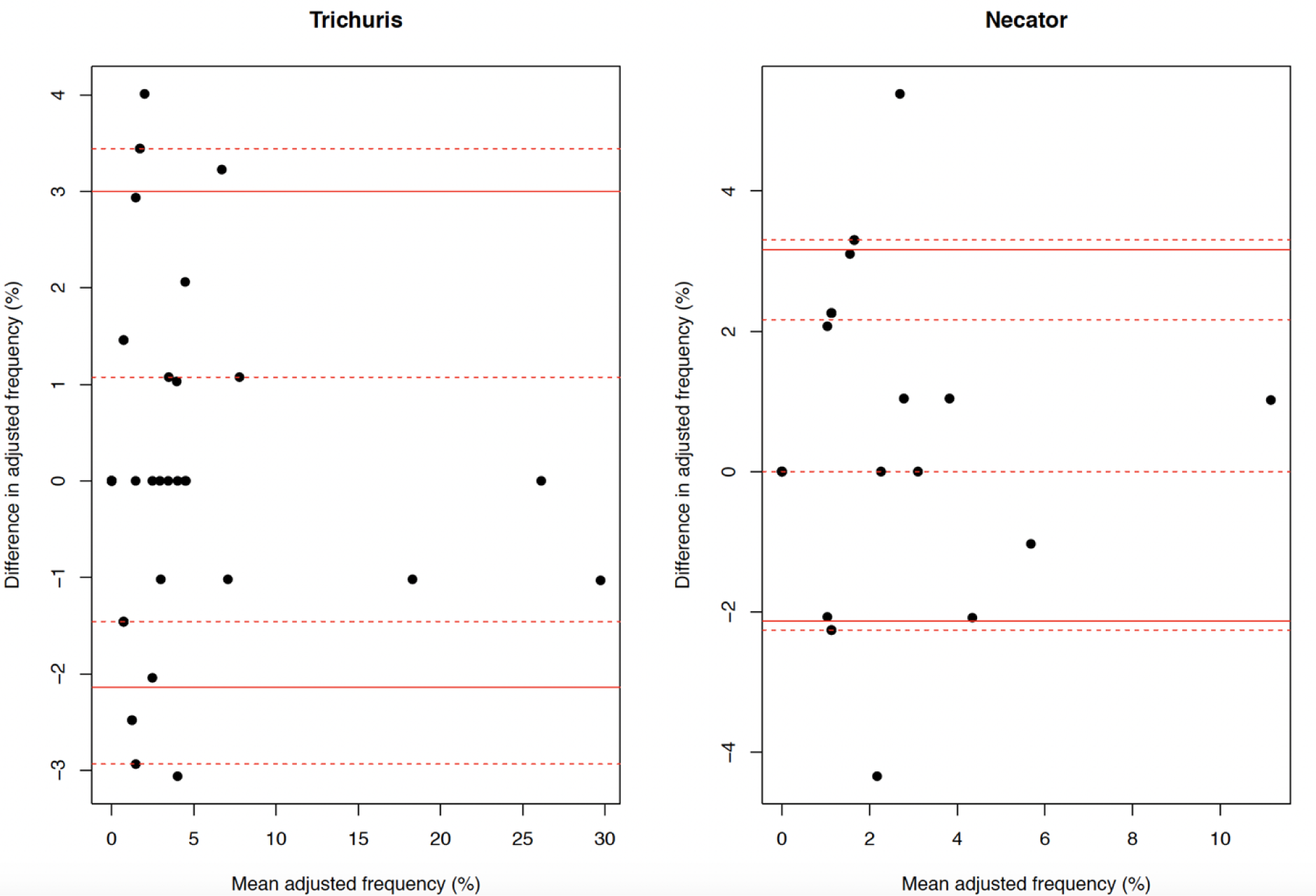
The Bland Altman plots illustrating the repeatability of the pyrosequencing assays on field samples for *Trichuris* and *Necator*. The straight line illustrates the change in adjusted frequency of mutant (MT) SNPs in the ß-tubulin isotype 1 for *Trichuris* (**top row**) and *Necator* (**bottom row**) for codons 167, 198 and 200 before (BL) and after (FU) administration of a single oral dose of 400 mg albendazole. The black lines represent subjects with a satisfactory response, while the red lines represent those that poorly or did not respond the treatment.

## Discussion

### Need to assess the analytical performance of assays that detect and quantify SNPs in ß-tubulin

To readily interpret the results obtained from the stool samples, we comprehensively evaluated the analytical performance of the pyrosequencing assays. For this, we applied the assays on both plasmid reference DNA with known MT:WT ratios (LoB, LoD, accuracy and precision) and stool samples (repeatability of test results). The outcome of this evaluation indicated that the LoB was non-zero (∼3.5%). This is not unexpected and in line with the technical limitations of the platform. It also underscores that the low frequency of MT SNP reported in previous studies should be interpret with caution as MT SNPs might actually be absent [16, 31, 33]. Other consequences of this suboptimal analytical specificity, are the relatively high LoD (2% – 7%). And the systematic bias towards higher MT SNPs (**Figs 1 – 3**). The relatively high LoD may question pyrosequencing as the preferred platform to accurately detect early signs of AR. This is in particular when the coefficient of variance exceeded 10% for samples with a low ratio of MT:WT. This trend of less precise test as a function of lower ratio of MT:WT was also confirmed by repeating the test results on stool samples (**Fig 5**).

This evaluation also revealed important differences in analytical performance across both codons and STH species, which further underscores the need of assessing and reporting the analytical performance of each individual assay. Moreover, to allow for comparing test results across studies, to identify the best platform/assay and for quality control, there is need for an external quality assurance scheme. During such a scheme test result that are independently reported by a laboratory are compared to those reported by reference or expert laboratories. This was recently piloted for DNA-based detection of STH, *Schistosoma* and *Strongyloidiasis* in stool, already revealing important differences across laboratories [44], which might be due to factors including but not limited to DNA-extraction method and PCR.

### Differences in occurrence and frequency of MT SNPs across both codons and STH are in line with literature

Except for codon 167 in *Ascaris* (we did not include this codon in any analysis for this STH), all MT SNP occurred in all codons and STHs (**Table 6**). The most prevalent MT SNPs were those in codon 200 for *Trichuris* (44.9% of the samples) and in codon 198 for *Necator* (12.6% of the samples), while the least prevalent MT SNP were both codons 198 (1.0% of the samples) and 200 (5.6% of the samples) of *Ascaris*. The frequency of MT SNPs within a sample was low. The highest frequency was observed in a *Trichuris* sample for which the adjusted MT SNP frequency at codon 167 equalled 26.1%. In the remaining samples the adjusted frequency of MT SNPs did not exceed 20%. Generally, these differences across both codons and STH are not unexpected and are very much in line with current scientific literature [16, 33, 34].

### Evaluation of evidence for the role of known SNPs in the ß-tubulin gene as markers for BZ resistance for the different human STHs

We assessed the role of the known SNPs in specific ß-tubulin genes as markers for BZ resistance based on trends across the following indicators: (i) drug pressure history and (ii) individual drug response, and (iii) change in both MT SNP occurrence and frequency following drug administration. Our results indicated that the SNPs matched the expectations for each of these three indicators and the researched STHs only for the following indications. For codon 198 in *Necator* and codon 200 in *Trichuris*, significant differences were noted, though not for every indicator. For codon 198 in *Necator*, the change in both the occurrence (proportion of samples) and the frequency of MT SNPs within samples followed a logical trend across the different study sites (proportion of samples containing MT SNPs / frequency of MT SNPs within a sample increasing from Ethiopia (short history of drug donation programs and ERR based on duplicate Kato-Katz: 96.3%; Laos PDR: moderate history of drug donation program and ERR: 96.2% and Pemba Island: long history of drug donation program and ERR: 83.6%). Significant differences in the occurrence and frequency of MT SNPs were also observed for codon 200 in *Trichuris* but the trend across study sites did not follow strict drug pressure history or therapeutic efficacy (Ethiopia: 48.1%, Lao PDR: 40.5%, Pemba Island: −4.9%). The adjusted frequency of MT SNPs increased following drug administration for *Trichuris* and *Necator*, though this change was significant for codon 200 of *Trichuris* only. This increase in MT SNPs following treatment was also reported by Diawara and colleagues [16] for *Trichuris* in codon 198 (isolates from Haiti) and codon 200 (isolates from Haiti and Kenya).

BZ resistance in veterinary nematodes can involve different β-tubulin SNPs in populations of the same parasite from different locations. For example, *Haemonchus contortus* which is a Clade V hematophagous roundworm similar to *Necator* but prevalent in sheep and other herbivores, exhibits a great deal of BZ resistance around the world. However, different locations can have different frequencies of different β-tubulin SNPs at codons 167, 198 and 200, causing that resistance [45]. In *Ancylostoma caninum*, a species even more closely related to *N. americanus*, BZ resistance has become a serious problem and is caused by the codon 167 mutation. Note that also another codon was identified (codon 134: CAA to CAT) [46]. However, BZ resistance has rarely been reported in Clade I and Clade III veterinary nematodes, such as *Trichuris* spp. and *Ascaris* spp. and genetic studies are lacking.

### Need for whole genome sequencing of well characterized worms

An important challenge to study BZ resistance in human STH is the access to phenotypic BZ resistant worms. While in veterinary medicine, phenotypic BZ resistant worms can be both easily obtained (under dosing of the animals and experimentally infecting animals with progeny of the surviving worms) and collected (animals can be slaughtered to collect worms *in-situ*), the best option to collect human phenotypic BZ resistant is probably limited to expulsion studies only. Moreover, other genes other than the ß-tubulin non-tubulin could play a role in BZ-resistance, highlighting the need for a wider analysis of the genome of intestinal worms. As part of the Starworms project, we conducted two expulsion studies in Pemba Island, with the aim to collect susceptible (worms that were expelled following a standard BZ treatment) and resilient *Trichuris* and hookworm populations (worms that survived the standard BZ treatment but were expelled after the administration of oxantel (*Trichuris*) or pyrantel (hookworms). We will conduct whole genome sequencing, after which a variance calling across susceptible and resilient worms will be performed.

### Sample preparation prior DNA extraction to maximize DNA yield needs attention

A major challenge during this study was the poor amplification rates. In approximately 25% of the samples, PCR was unsuccessful, even if we repeated the PCR using the product of the first PCR as a template. It is well known that extracting DNA out of STH eggs in stool samples is not trivial [47]. This is particularly true for *Trichuris* and *Ascaris*, for which DNA is well-protected by a thick eggshell (*Necator* eggs have a thin shell). This variation in amplification rates across STH species is also apparent in our study, amplification rates ranging from 52.9 – 60.9% for *Trichuris*, 75.6 – 77.4% for *Ascaris* and 67.6 – 81.2% for *Necator*. Remarkable is that each of the DNA extracts resulted in a positive species-specific qPCR, though on a multicopy gene [48]. This does not only further explain the relatively lower amplification rates, it also has important implications on researching the MT SNPs ß-tubulin genes as markers for BZ resistance in human STH. First, information on the MT SNPs is not available in a considerable proportion of the samples. Second, the frequency of MT SNP might be based on a low number of copies of ß-tubulin, and hence may further impede a readily interpretation of the observed pyrosequencing results. This is in particular for follow-up samples where the egg counts will be already low. To maximize the DNA yield, it will therefore be important to research and standardize alternative sampling processes. In veterinary medicine, DNA yield of animal STH is sometimes increased by culturing samples and subsequently recovering larval stages [49]. These larval do not only consist of multiple cells (*vs.* single cells in STH egg), they are also hatched (DNA is not protected by any eggs shell) resulting in a satisfactory DNA yield. Although this procedure could be applied for human hookworms applying the Harada Mori technique [50], it does not apply for *Ascaris* and *Trichuris*. For these STHs, the larval stages do not hatch outside of the host. An alternative is to increase the number of eggs subjected to DNA extraction by purifying eggs from a larger quantity of stool. Pooling stool samples of multiple individuals might also be considered. This will not only increase the number of eggs, it also a cost-saving strategy allowing the screening of MT SNPs at a larger scale. Such an approach has been used in veterinary medicine to screen the occurrence of MT SNPs across sheep flocks across the UK [51]. Finally, one could also further improve the sensitivity of the assay (e.g., different polymerases, master mixes)

### Need for absolute quantification of SNPs

While the pyrosequencing results provide a ratio of MT SNPs only, it will be important to explore methods that allow for absolute quantification of SNPs too. This is in particular when DNA-yields are low. For example, it is expected that a gene copy ratio of 1:10 MT:WT or 10:100 MT:WT would result in both scenarios both a 90% MT, yet it is clear that in the second scenario the estimated proportion will be more reliable. Platforms that allow for absolute quantification are droplet PCR (dPCR) protocols to quantify MT SNPs in animal STH have been developed [52].

## Conclusions

Our study underscores the lack of compelling evidence for the role of the known SNPs in the ß-tubulin genes examined, as markers for benzimidazole resistance in the different STH species. It also highlights that there is a need to assess the analytical performance of any assays in order to readily interpret and compare results. Further research should therefore also focus on genes other than the ß-tubulin genes and/or on whole genome sequencing and comparison of putative resistant and susceptible STH isolates/samples, if and when such characterized samples become available.

## Supplementary Information

**Table S1.**
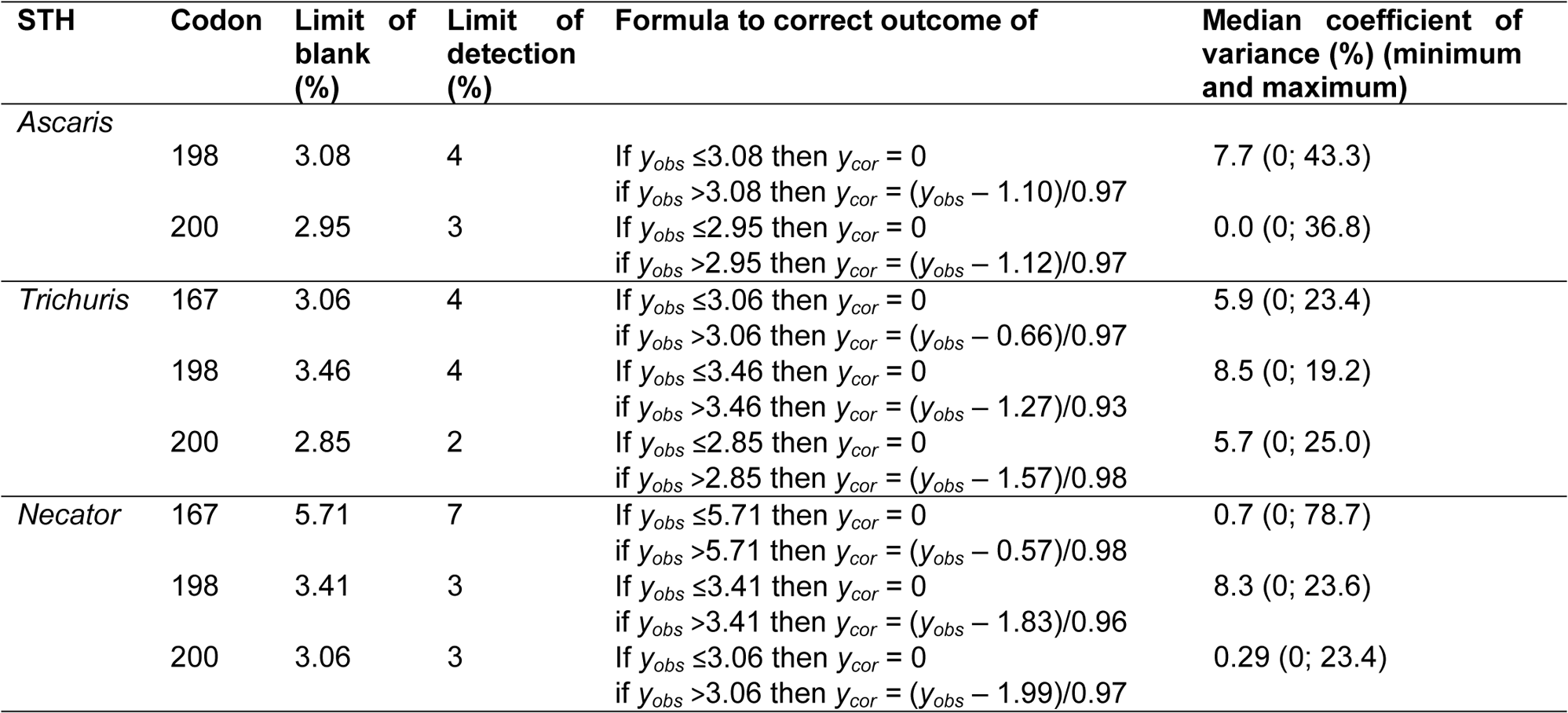
The diagnostic performance of the pyrosequencing assay for the detection and quantification of single nucleotide polymorphisms.

**Info S1. The data set used to determine the analytical performance of the pyrosequencing assays and the occurrence of SNPs in ß-tubulin in stool samples.**

